# Medial prefrontal cortex has a causal role in selectively enhanced consolidation of emotional memories after a 24-hour delay: A TBS study

**DOI:** 10.1101/2020.10.11.335125

**Authors:** Nicholas Yeh, Jessica D. Payne, Sara Y. Kim, Elizabeth A. Kensinger, Joshua D. Koen, Nathan S. Rose

## Abstract

Previous research points to an association between retrieval-related activity in the medial prefrontal cortex (mPFC) and preservation of emotional information compared to co-occurring neutral information following sleep. Although the role of the mPFC in emotional memory likely begins at encoding, little research has examined how mPFC activity during encoding interacts with consolidation processes to enhance emotional memory. This issue was addressed in the present study using transcranial magnetic stimulation in conjunction with an emotional memory paradigm. Healthy young adults encoded negative and neutral scenes while undergoing concurrent TMS with a modified short intermittent theta burst stimulation (sTBS) protocol. Participants received stimulation to either the mPFC or an active control site (motor cortex) during the encoding phase. Recognition memory for scene components (objects and backgrounds) was assessed after a short (30-minute) and a long delay (24-hour, including a night of sleep) to obtain measures of specific and gist-based memory processes. The results demonstrated that, relative to control stimulation, sTBS to the mPFC enhanced memory for negative objects on the long delay test (collapsed across specific and gist-based memory measures). mPFC stimulation had no discernable effect on memory for objects on the short delay test nor on the background images at either test. These results suggest that mPFC activity occurring during encoding interacts with consolidation processes to preferentially preserve negatively salient information.

**Significance Statement:** Understanding how emotional information is remembered over time is critical to understanding memory in the real world. The present study used noninvasive brain stimulation (repetitive transcranial magnetic stimulation, rTMS) to investigate the interplay between mPFC activity that occurs during memory encoding and its subsequent interactions with consolidation processes. rTMS delivered to the mPFC during encoding enhanced memory for negatively valenced pictures on a test following a 24-hr delay, with no such effect on a test occurring shortly after the encoding phase. These results are consistent with the hypothesis that emotional aspects of memories are differentially subjected to consolidation processes, and that the mPFC might contribute to this “tag-and-capture” mechanism during the initial formation of such memories.

## Introduction

Emotional memory is often characterized by “trade-off effects”, where superior recollection of the emotional parts of an experience occur at the expense of memory for neutral aspects (Kensinger et al., 2007a; Payne et al., 2008). One account of emotional memory trade-offs proposes that synaptic processes operating near the initial learning event can “tag” emotionally salient aspects of an experience and set the stage for downstream preferential consolidation of these emotional aspects (Kim & Payne, 2020; Payne & Kensinger, 2018; also see, Richter-Levin & Akirav, 2003). This “tag and capture” mechanism has been proposed to explain how encoding and subsequent consolidation processes interact to promote later stabilization of memories. While the synaptic tag has yet to be directly examined in humans, recent studies have observed network-level neural processes via functional MRI (Tambini et al., 2017) and behavioral outcomes (Ballarini et al., 2013; Dunsmoor et al., 2015; Patil et al., 2016) that are consistent with the theorized tagging mechanism.

An open question in memory research is whether these network-level neural processes can be utilized to causally manipulate the preferential encoding and consolidation of emotional information. Neuroimaging studies have established that increased levels of activation of the medial temporal lobe (MTL; including the amygdala and hippocampus) and the medial prefrontal cortex (mPFC) are associated with successful emotional encoding of negative information (cf. Murty et al., 2010; Payne & Kensinger, 2010, 2011, 2018; see also Goto & Grace, 2008). Functional connectivity between the MTL and mPFC is also associated with successful emotional memory encoding (Berkers et al., 2016; Kensinger & Corkin, 2004).

There is also evidence that longer consolidation delays (e.g., 24-hour delays) are crucial for emotion-related memory enhancements (Dunsmoor et al., 2015; see also Patil et al., 2016 for reward-related memory). One probable reason for this delay-dependency is the presence of sleep during the consolidation interval. Sleep-based consolidation processes play a pivotal role in emotional memory trade-off effects by selectively preserving negative information (Payne et al., 2008), especially gist-based aspects of negative information (Payne et al., 2008; Hu et al., 2006). Evidence from fMRI studies suggests that sleep leads to a refinement in the neural networks engaged in emotional memory retrieval. For example, studies have revealed that initial broad network activation observed during encoding is refined into a smaller network centered on the amygdala, hippocampus, and ventral mPFC following a sleep-filled consolidation delay, with greater connectivity among these regions correlating with enhanced memory for negative information compared to wakefulness (Bennion et al., 2015; Payne & Kensinger, 2011; Sterpenich et al., 2009). This refinement in emotional memory retrieval networks presumably occurs as memories are consolidated.

Importantly, the hypothesized tag-and-capture mechanisms that lead to refinements in emotional memory retrieval networks are established near the time of encoding. Payne and Kensinger (2018) proposed that successful emotional memory will be optimal with increased MTL and PFC activity near encoding, with sleep occurring shortly after. Critically, emotional tags can be set via stress- and arousal-related neuromodulators that reflect initial encoding activity between the MTL and PFC. However, sleep is required to ensure that these tags persist (Payne & Kensinger, 2018), suggesting that post-learning sleep is essential for transforming temporary synaptic changes into long-lasting systems-level ones, and for linking these distributed tags into an integrated memory trace.

Thus, by modulating neural activity at encoding, we may be able to affect how emotional information is subsequently consolidated. The present work therefore sought to understand how the mPFC might influence interactions between encoding and consolidation processes that are thought to lead to selective preservation of negative memories (Payne & Kensinger, 2018). During the encoding phase of an emotional memory trade-off task, we applied a short, modified version of intermittent theta burst stimulation (sTBS) to either the mPFC or an active control site (motor cortex; MC) (see the “Transcranial Magnetic Stimulation” section in the methods for details about the sTBS protocol and the rationale for how we targeted the mPFC). A recognition test assessed memory following a short (30-minute) delay and a long (24-hour) delay that included sleep.

Under the assumption that mPFC activity at encoding interacts with subsequent consolidation processes to refine the MTL-mPFC network associated with memory for negative information, we assumed that modulating mPFC with sTBS during an emotional memory encoding task would, in turn, alter activity in an excitatory manner to both the targeted and functionally connected areas (similar to the effects observed with standard intermittent TBS, Hermiller et al., 2020; Huang et al., 2005; Tang et al., 2019) within an emotion–cognition network, including changes in MTL-mPFC connectivity. Enhanced emotional memory performance at the short delay would be suggestive of differences in encoding (and very early consolidation processes) due to mPFC stimulation, while altered behavioral outcomes at the long delay, especially in the absence of differences at the short delay, would indicate that TMS-related modulation of this network prioritized certain items during encoding for *later* consolidation (e.g., post-sleep). Our *a priori* hypothesis was that mPFC relative to control stimulation would preferentially enhance memory for negative, but not neutral, information following a long delay. Furthermore, this study also examined the effects of mPFC stimulation during encoding on the specificity of these emotional memory trade-off effects for specific and gist-like emotional memories (cf. Bovy et al., 2020; Hu et al., 2006; Payne et al., 2008).

## Materials and Methods

### Participants

Forty-five participants (aged 18-24; *M* = 19.71; N_females_= 30) were recruited from the University of Notre Dame. Participants identified as 71% Caucasian, 11% Asian, 9% African American, 7% Hispanic, and 2% who declined to state. All participants were fluent English speakers, had no history of sleep disturbances, and did not take medication known to impact sleep or present contraindications for TMS. The current sample size was selected to be in line with prior TMS work (Sack et al., 2009) that has detected significant behavioral effects when targeting cortical regions based on the 10-20 system (see TMS section below). Participants were randomly assigned to receive sTBS either to the mPFC (*N* = 23) or to motor cortex (MC; *N* = 22), which served as an active control site.

Participants were screened for contraindications for TMS following published criteria (Rossi et al., 2009), and were excluded if they self-reported any history of seizures, brain injuries (e.g., stroke, aneurysms, concussions, traumatic brain injury), severe headaches/migraines, fainting, metal implants, neurological/psychiatric disorders, potential pregnancy, or psychoactive medication use. Additionally, participants were instructed to get no less than six hours of sleep the night before and refrain from caffeine, alcohol, and tobacco three hours prior to the study. Upon arrival, participants provided informed consent. Following completion of the study, they received either course credit or monetary compensation ($15/hour).

### Materials

The stimuli have been used in previous work (Payne et al., 2008) with 448 total critical scene components (objects and backgrounds) selected in this study. The critical scene components were then partitioned into old (256) and new items (192). The encoding task consisted of 128 complex valenced scenes (64 negative, 64 neutral) that were created by placing a negative (e.g., snake) or neutral object (e.g., chipmunk) on a plausible neutral background (e.g., forest). The short and long delayed recognition tests each contained a unique set of old and new scene components (i.e., objects and background in isolation). Studied scenes consisted of the objects and the background from “whole” scenes, i.e., objects on backgrounds that were presented together during the study phase. New scene components were drawn from scenes that were not presented during the study phase. Each recognition test consisted of 224 total scene components: 32 previously viewed (“same”) objects (16 negative, 16 neutral), 32 similar objects (16 negative, 16 neutral), 64 foil objects (32 negative, 32 neutral), 32 same backgrounds (16 shown with negative object, 16 shown with a neutral object), 32 similar backgrounds (16 similar objects were shown with a negative object, and 16 with a neutral object), and 32 foil backgrounds (all neutral). Same components were defined as the exact same background or object studied during encoding (e.g., identical snake). Similar components were defined as an alternative version of a background or object that differed in any number of visual features from those studied during encoding (e.g., a snake that differed in color, positioning, shape, species, etc.). Note, since backgrounds were always non-emotional, there are no negative backgrounds. The scene components were never presented twice (i.e., in both tests) and were counterbalanced. Additionally, participants never saw both a same (e.g., same snake) and similar (e.g., similar snake) scene during the one test.

### Experimental design

Participants completed the experiment over the course of two sessions separated by approximately 24-hours. The first session included determination of active motor threshold (see section Transcranial Magnetic Stimulation below), the encoding task, and a recognition test after a 30-minute delay (“short delay” test). sTBS was administered during the encoding phase with a 30-minute delay occurring from the completion of the encoding phase to the onset of the recognition test. Note that simultaneous electroencephalography recordings were obtained during the encoding and short-delay recognition tasks, but not in the final (24-hour) delayed recognition test. These data are not the focus of the present manuscript and therefore will not be discussed further. After the first session, participants returned to the lab approximately 24-hours later to complete a recognition test on the remaining untested items (“long delay” test).

### Emotional Memory Trade-off Task

#### Encoding Phase (Figure 1a)

Participants were instructed that they would be presented with a series of scenes and to imagine that they were coming across the scenes in real life. Participants viewed 128 negative and neutral complex scenes. For each scene, they made an approach/avoid judgment using a 7-point scale (1 = move closer, 7 = move away) to facilitate incidental encoding. The scenes were presented pseudorandomized in 16 blocks with 8 images per block (4 negative, 4 neutral). Each trial began with a 100-200ms jitter prior to a 1000ms fixation cross. Following the fixation cross, a valenced scene was presented for 3000ms and was replaced with a self-paced avoid/approach judgment scale. Following the 8^th^ trial of each block, participants underwent 2s of sTBS (30 pulses), for a total of 480 pulses over the entire encoding phase.

**Figure 1.**
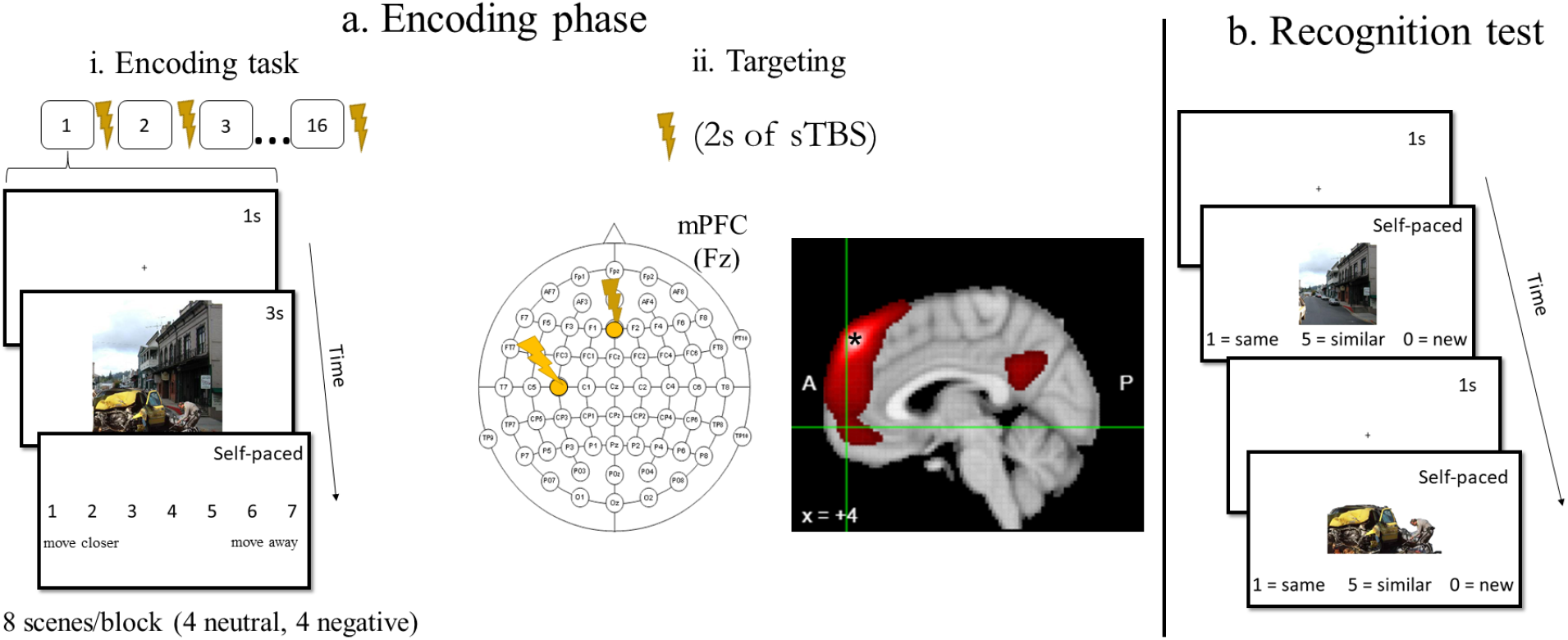
(a) Encoding phase. (i) Encoding trial structure and timing. Participants viewed a fixation cross (1s), followed by a valenced image (3s) and an approach/avoid scale (self-paced). Stimulation occurred after every 8^th^ image for a total of 128 images and 16 blocks. Participants underwent stimulation to only one site, which was randomly assigned. (ii) The Fz and C3 electrodes were targeted for mPFC (N = 23) and motor control stimulation (N = 22), respectively. The Fz electrode was selected based on its proximity to the dorsal mPFC region (MNI: 4, 52, 46; indicated by *) that is functionally connected with the ventral mPFC region (MNI; 4, 56, −8; indicated by the green crosshair) that was linked to the selective preservation of negative objects in memory from Payne and Kensinger (2011). We targeted the dmPFC (Fz) under the assumption that sTBS would modulate neural activity in an excitatory manner (similar to iTBS) in both the target site and its associated networks. (b) Recognition task. Same, similar, and new objects (negative and neutral) and backgrounds (neutral) were presented individually. Participants responded “same”, “similar”, or “new”. Scene components were presented in randomized order.

#### Recognition Memory Test

Participants were instructed to make same, similar, or new judgments to old and new scene components on both the short and long delayed recognition tests. Participants completed a short practice phase to familiarize themselves with the recognition test and the same, similar, and new judgments before the recognition test (see Figure 1b). Same judgments were defined as the exact same background or object studied during encoding (e.g., identical snake). Similar judgments were defined as an alternative version of a background or object that differed in a specific visual detail from those studied during encoding (e.g., a snake that differed in color, positioning, species, etc.).

Each trial began with a 100-200ms (10ms steps) jittered interval prior to a 1000ms fixation cross. Following the fixation cross, participants made self-paced same, similar, or new judgments of individually presented negative objects, neutral objects, or neutral backgrounds. Objects and backgrounds were presented randomly, but only once, in either the short or long delay recognition test.

### Transcranial magnetic stimulation

A PowerMag EEG 100 TMS stimulator (Mag & More GmbH., Munich, Germany) and a 70 mm figure-eight coil (PMD70-pCool) were used for administration of TMS. Active motor threshold was obtained by placing the coil over the left motor cortex (near the C3 electrode) and locating the site that produced visible movement in the right thumb from single pulses of TMS. Active motor threshold was defined as the lowest percentage of stimulator output that elicited visible movement in the right thumb on 5 out of 10 trials while participants maintained contraction of the right thumb and index finger. Visible muscle twitches were determined by the first author in each session. This was done to obviate variability in what constitutes a visible muscle twitch across sessions and experiments. We determined active motor threshold and sTBS intensity while participants were wearing the EEG cap, so the coil distance from the stimulation site was the same during active motor thresholding and the encoding task sTBS. The mean motor threshold was matched between participants receiving mPFC and MC stimulation (mPFC: *M* = 59%, *SD* = 10%; MC: *M* = 59%, *SD* = 11%; *t*(42) = −.25, *p* = .805). As is standard, the sTBS protocol was administered at 80% of active motor threshold.

The stimulation site for the dorsal mPFC was determined using a large-scale meta-analysis of 340 fMRI datasets using Neurosynth.org (Yarkoni et al., 2011). The Fz electrode was selected as the dorsal mPFC stimulation site based on the Neurosynth meta-analysis that showed the MNI coordinate under the Fz electrode (MNI: 4, 52, 46; Okamoto et al., 2004; Okamoto & Dan, 2005) had strong functional connectivity and coactivation with the ventral mPFC site identified by Payne and Kensinger (2011) that was associated with enhanced negative object recognition following sleep compared to wakefulness (MNI = 4, 56, −8; Figure 1b). The rationale for targeting the dorsal mPFC is that TMS modulates neural activity in both the target site and its associated networks (for an example with TMS to cortical regions modulating the hippocampus, see Wang et al., 2014; Warren et al., 2019). Thus, we reasoned that stimulating the dorsal mPFC would also affect the ventral mPFC activity, which is not directly accessible to TMS. For the control stimulation site, we targeted the MC under the C3 electrode because it is both a common control site (e.g., Daskalakis et al., 2008; Fecchio et al., 2017) and, importantly, we are not aware of any evidence demonstrating a strong connectivity between the MC and the ventral mPFC region during emotional encoding.

The typical offline protocol for iTBS follows 2s on/8s off cycles that deliver 600 pulses with each 2s of stimulation corresponding to 3 pulses at 50Hz with an inter stimulus interval of 200ms. We adopted a slightly modified sTBS approach with identical 2s periods of stimulation that was applied in a similar 2s on/8s off manner that was interleaved (between each encoding block) throughout the encoding phase. Specifically, following each encoding block (8 images), participants were instructed to close their eyes and relax while the TMS coil was positioned to either the mPFC or MC stimulation site location. For the mPFC condition, the coil was positioned at 0° from the midline pointing posteriorly to target the Fz electrode. For the MC condition, the coil was positioned at 45° from the sagittal plane to target the C3 electrode (Figure 1b). Following 2s of sTBS, participants were instructed to continue with the encoding task with a self-paced button press. The average time from the last TMS pulse to the onset of the first and last image in the following encoding block was 8.28 seconds (range: 6.14 - 19.66) and 48.71 seconds (range: 45.62 - 60.01), respectively.

A similar approach has previously been shown to facilitate memory and, importantly, to not have cumulative effects on performance (Demeter et al., 2016). Prior to the start of the experimental task, participants received one 2s train of sTBS to become familiarized with the procedure. No participants reported any adverse response to sTBS.

### Statistical analysis

For the recognition data, participants’ memory scores were calculated for general recognition (gist-familiarity) and specific recognition (recollection). The contrast between general and specific recognition scores was our focus because prior research suggests that overnight sleep exerts its strongest effects on gist-based familiarity, rather than specific recollection of memory details (Payne et al., 2008). In line with prior studies using a same-similar-new judgment at retrieval (Garoff et al., 2005; Kensinger et al., 2007b), general recognition parallels the independence-formulas score commonly used in Remember/Know paradigms (Yonelinas & Jacoby, 1995) with the following formula:

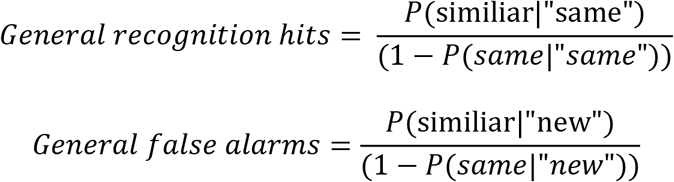

Thus, general hits are calculated based on when participants respond “similar” to previously viewed items, taking into account when participants respond “same” to previously viewed items. General false alarms are measured when participants respond “similar” to new items, taking into account when they respond “same” to new items. Specific recognition was scored with the following formula:

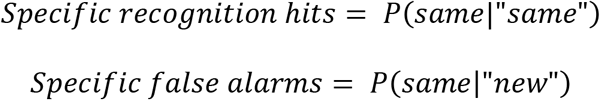

Thus, specific hits and false alarms are computed when participants respond “same” to previously viewed items and “same” to new items, respectively.

Since our primary aim was to investigate the effects of mPFC sTBS on emotional memory we computed recognitions scores for negative objects, neutral objects, the neutral backgrounds that were originally paired with a negative object, and the neutral backgrounds that were originally paired with a neutral object. Importantly, these calculations were repeated for the short and long delay tests. This enabled us to systematically compare how mPFC versus MC sTBS affected memory for negative and neutral objects, backgrounds in negative and neutral scenes, across the short and long delay tests. Analyses focused on memory performance for “same”, “similar”, and “new” responses to the same and new scene components (i.e., responses to similar objects and background scenes were not analyzed), to be consistent with previous work using this paradigm (Kensinger et al., 2007a, 2007b; Waring et al., 2010).

There were statistically comparable ‘same’ (range = .038-.096) and ‘similar’ (range = .153-.258) false alarm rates between the cells in the experiment (all *p*’s > .05), which is consistent with previous work (Payne et al., 2008). All memory performance analyses were conducted on corrected recognition scores (hits – false alarms) to obviate issues with response bias effects. We will refer to these measures as recognition scores from this point forward.

Analyses were conducted in R (version 3.6.3.) with the *afex* (Singmann et al., 2020) and *emmeans* packages (Lenth, 2020). Significant effects in the ANOVAs were followed-up with post-hoc contrasts as implemented in the *emmeans* package. Degrees of freedom for post-hoc contrasts were estimated using the Kenward-Rogers method. Three participants (mPFC: 2, MC: 1) were excluded from the analysis due to no observations in one or more response categories for the general recognition test. Results are considered significant at an alpha level of .05 unless otherwise noted and post hoc degrees of freedom were estimated with the Kenward-Roger method.

## Results

We tested our *a priori* hypothesis that, relative to MC stimulation, mPFC stimulation would facilitate recognition for negative objects at a long delay using a 2 (stimulation site: mPFC, MC) × 2 (valence: negative, neutral) × 2 (scene component: object, background) × 2 (delay: short, long) × 2 (Memory score: general, specific) ANOVA. The analysis included a between-subjects factor (stimulation site) and within-subject factors (valence, scene component, delay, and memory score). The full results are reported in Table 1, and the primary results of interest are reported below.

**Table 1.**
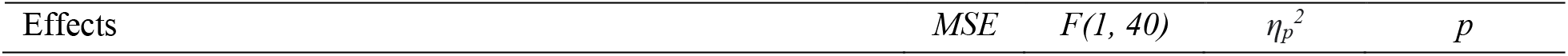
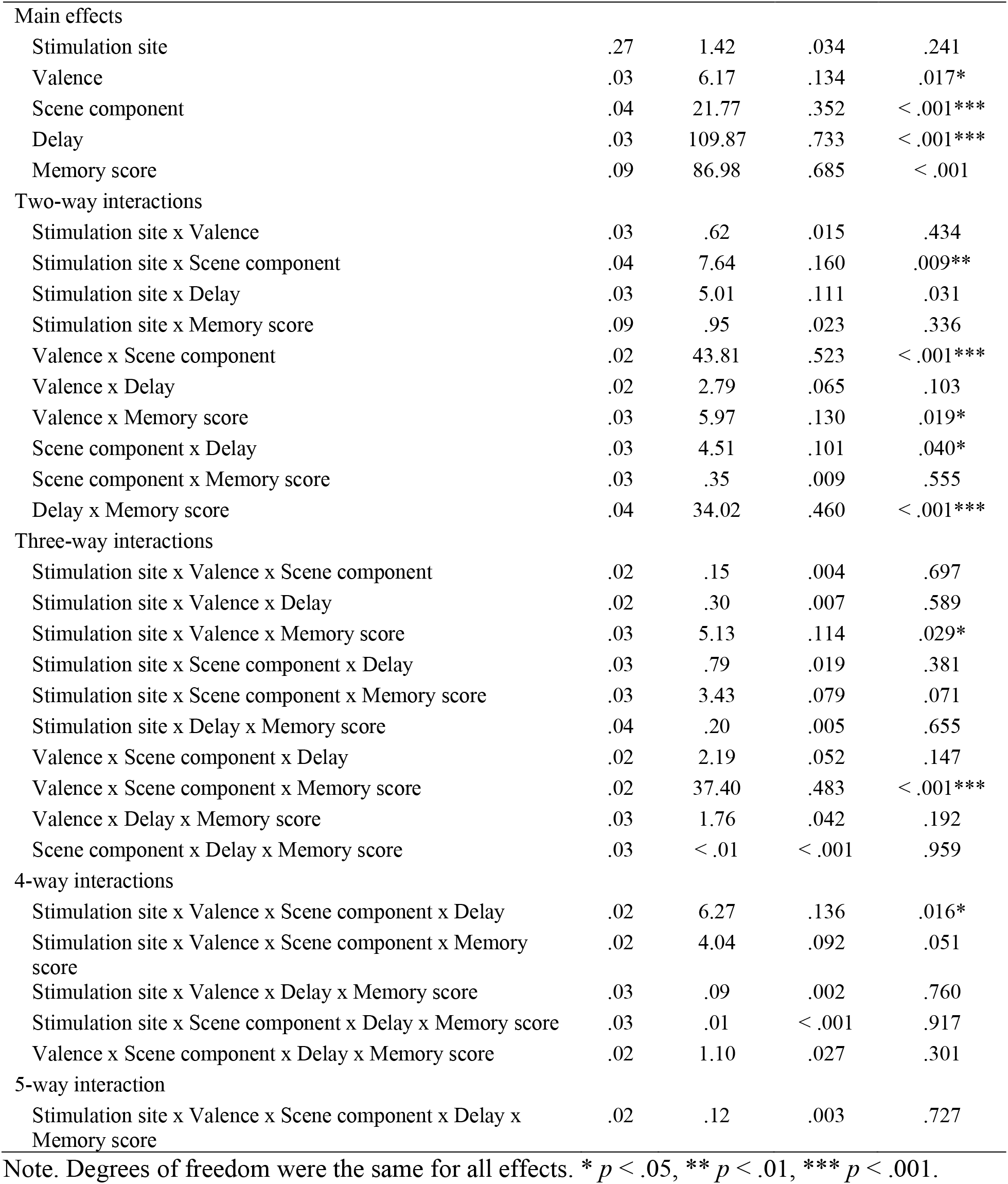
ANOVA results

### The emotional memory trade-off effect

In line with prior work (e.g., Payne et al., 2008), we first examined the existence of the emotional trade-off effect and if this effect remained consistent over time. As expected, we replicated the emotional memory trade-off effect as our results revealed a significant valence × scene component interaction, *F*(1, 40) = 43.81 *MSE* = .02, *p* < .011, *η_p_*^2^ = .523 (*Figure 2*). Follow up *t*-tests revealed that for negative scenes, objects were better remembered than their accompanying backgrounds, *t*(73.2) = 7.67, *p* < .001, *d* = .705. Critically, this pattern was not observed for neutral scenes, as memory performance did not differ between objects and backgrounds, *t*(73.2) = .14, *p* = .890, *d* = .013. Somewhat surprisingly, the scene component × valence × delay interaction revealed no evidence that the emotional memory trade-off differed between a short and long delay, *F*(1, 40) = 2.19, *MSE* = .02, *p* = .147, *η_p_*^2^ = .052.

**Figure 2.**
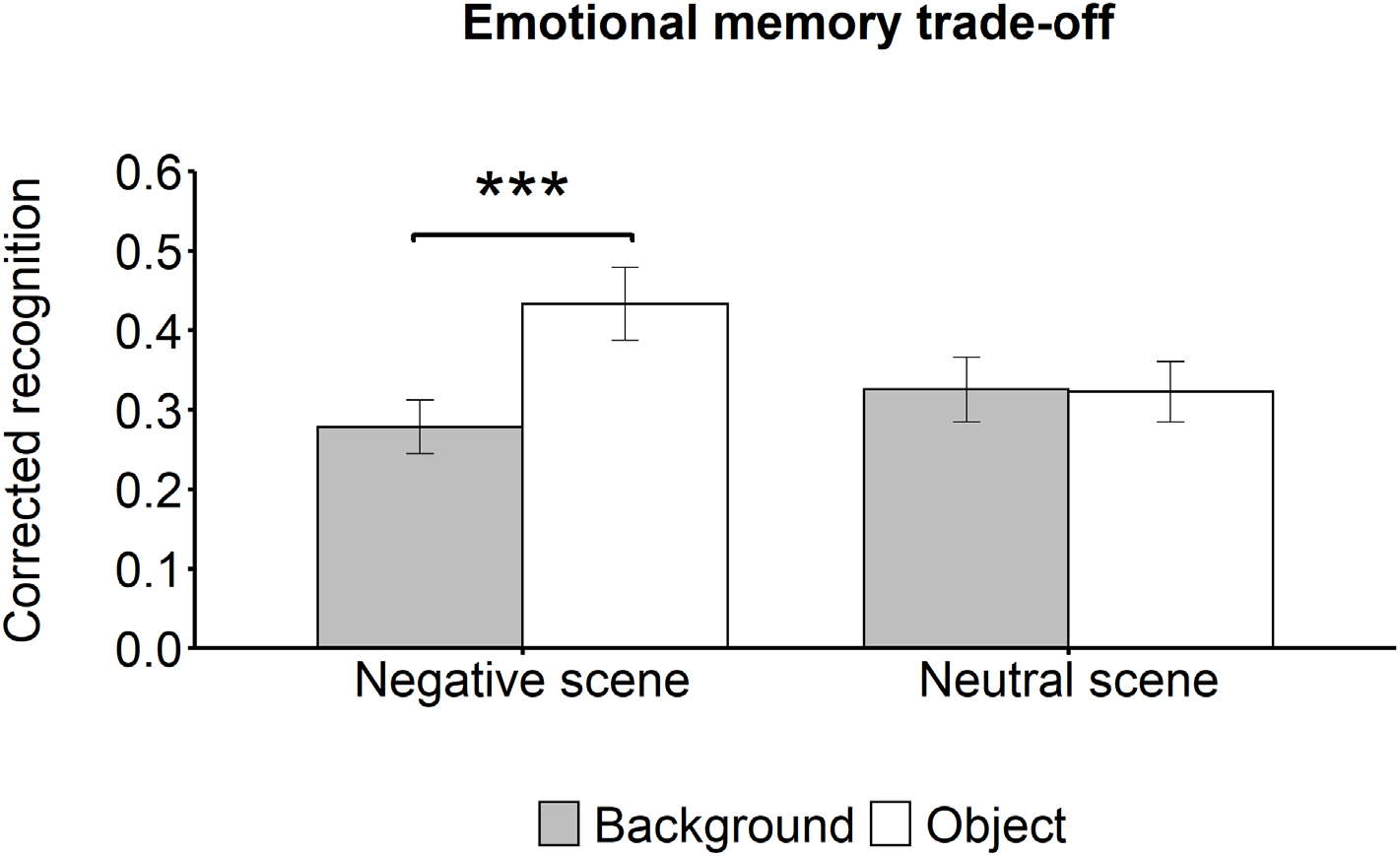
Corrected recognition scores (hits – false alarms) revealed an emotional memory trade-off effect with greater memory for objects compared to backgrounds in negative scenes. Additionally, no memory differences emerged for neutral scenes. Error bars represent 95% confidence intervals. \*\*\**p* < .001. Note, this is collapsed across stimulation site, delay, and memory scores.

### mPFC activity modulates the emotional memory trade-off effect at a long delay

We then examined the causal role of mPFC activity near the time of encoding interacting with consolidation processes (e.g., modulating the emotional memory trade-off effect). Because prior work has linked the mPFC in selectively preserving negative information following sleep, we anticipated that negative objects would be preferentially remembered following a long delay that included sleep in the mPFC condition relative to the MC condition. In line with our prediction, our analysis revealed a significant stimulation site × scene component × valence × delay interaction, which suggests that mPFC compared to MC sTBS modulated the emotional memory trade-off effect, *F*(1, 40) = 6.27, *MSE* = .02, *p* = .016, *η_p_*^2^ = .136 (*Figure 3)*.

**Figure 3.**
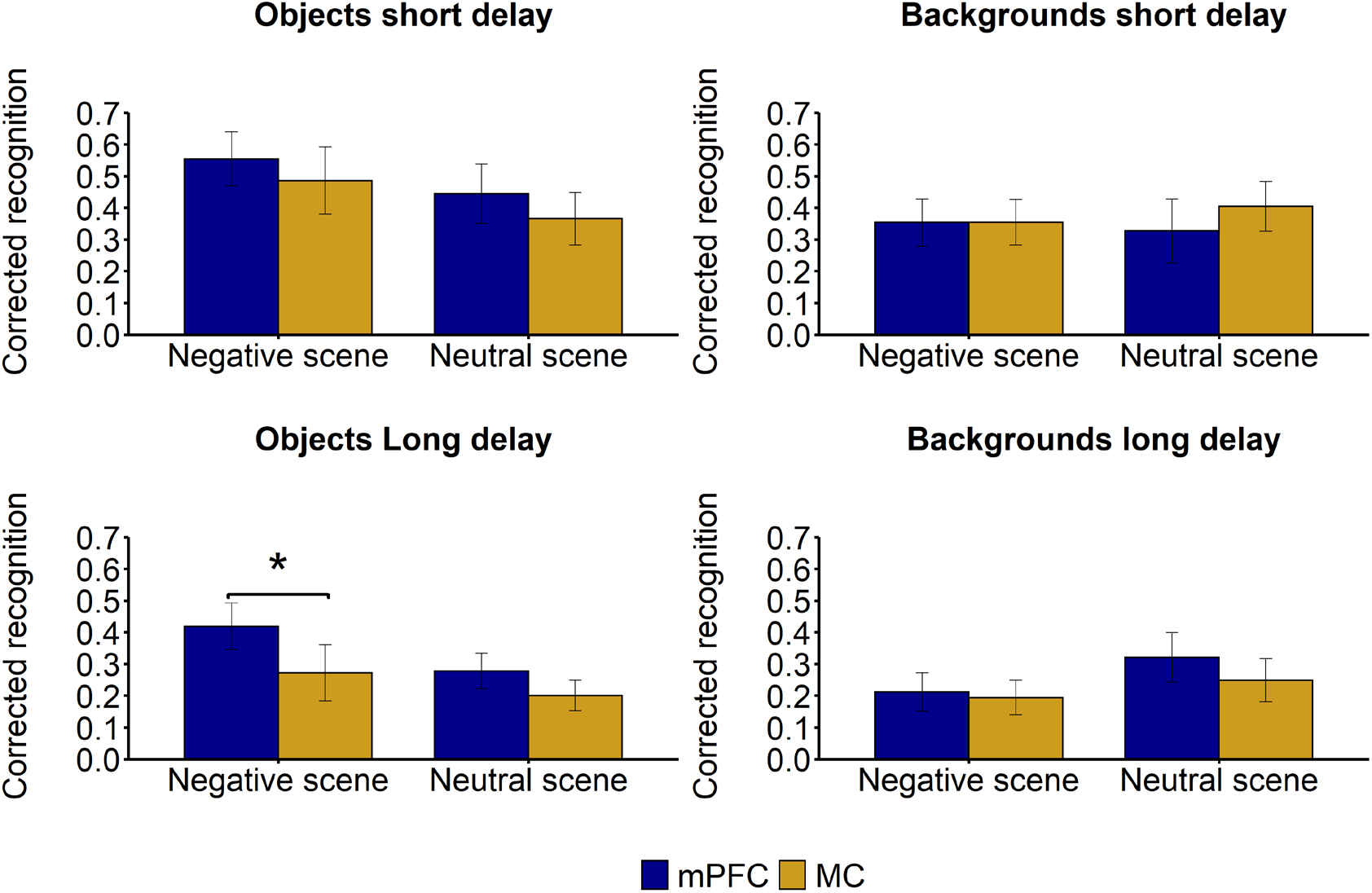
Corrected recognition scores (collapsed across memory score) revealed that mPFC compared to MC sTBS facilitated memory for negative objects at a long delay. No memory differences were found between negative objects at a short delay for mPFC vs. MC control stimulation. Error bars represent 95% confidence intervals. \**p* < .05.

To parse out the 4 way-interaction, follow up *t*-tests compared memory differences between the mPFC vs. MC stimulation conditions at each of the other factors (scene component, valence, delay). At the short delay, we observed no memory differences between the mPFC and MC stimulation site conditions for negative objects, *t*(110) = 1.30, *p* = .196, *d* = .313, or their backgrounds, *t*(110) = −.02, *p* = .988, *d* = −.004. Similarly, no memory differences emerged between the mPFC and MC stimulation site condition for neutral objects, *t*(110) = 1.49, *p* = .139, *d* = .358, or backgrounds, *t*(110) = −1.46, *p* = .147, *d* = −.351. Thus, we found no evidence that mPFC sTBS modulated the emotional memory trade-off effect at a short delay that did not include sleep (collapsed across memory score).

In line with our predictions, at a long delay, mPFC compared to MC stimulation facilitated memory for negative objects, *t*(110) = 2.78, *p* = .006, *d* = .669, with no differences emerging for their backgrounds, *t*(110) = .33, *p* = .739, *d* = .081. Importantly, we found no evidence for memory differences for neutral objects, *t*(110) = 1.47, *p* = .145, *d* = .353, or backgrounds, *t*(110) = 1.36, *p* = .175, *d* = .328. This pattern of results suggests that sTBS to the mPFC (compared to the MC) modulated the emotional memory trade-off effect by selectively preserving memory for negative objects, but not backgrounds, following a long delay that included a night of sleep (collapsed across specific and general memory scores). This effect was only found at the long delay, as no differences emerged for objects or backgrounds when memory was assessed at the short delay. Importantly, for neutral scenes, no differences between stimulation conditions emerged for objects or backgrounds when memory was assessed at either a short or long delay.

### The role of mPFC activity in memory specificity

Next, we sought to address if mPFC stimulation differentially modulated the emotional memory trade-off effect at a short or long delay when assessed on specific vs. general memory. The 5 way interaction found no evidence that mPFC vs. MC stimulation at a short or long delay differentially modulated the emotional memory trade-off effect when memory was assessed on specific and general recognition scores, *F*(1, 40) = .12, *MSE* = .02, *p* = .727, *η_p_*^2^ = .003. Lastly, we examined if mPFC sTBS differentially modulated the emotional memory trade-off effect collapsed across delay when corrected recognition scores were assessed on specific vs. general information (gist). Our results revealed a marginally significant 4-way interaction among stimulation site, valence, scene component, and memory type, *F*(1, 40) = 4.04, *MSE* = .02, *p* = .051, *η_p_*^2^ = .092. Although this did not reach significance, there appeared to be a numeric trend such that mPFC stimulation modulated the emotional memory trade-off effect for specific vs. general recognition (collapsed across short and long delays).

### Summary of Results

Collectively, the findings revealed that sTBS to the mPFC selectively facilitated recognition of negative information after a long delay while no differences emerged for their accompanying backgrounds. Critically, no memory differences emerged when memory was assessed following a short delay in negative scenes or at either a short or long delay in neutral scenes. Together, these findings suggest that modulating mPFC activity during encoding selectively facilitated emotional memories.

## General Discussion

This study investigated the causal role of the mPFC in the preferential encoding and consolidation of emotionally salient information. Our results provide preliminary evidence that the mPFC is causally involved in the encoding of negatively valenced scenes. Specifically, stimulating the mPFC, but not the MC, during encoding with sTBS enhanced memory for negative objects only after a 24-hour delay (filled with sleep). Moreover, sTBS to the mPFC had no detectable effect on memory for negative stimuli on the short delay (30 minute) test, on neutral information on either test, or on the background scene component in any condition. Notably, false alarm rates did not differ between any of the conditions, which provides evidence that our results are due to effects on recognition, rather than differences in response bias between the mPFC and MC groups. We interpret these findings as being consistent with the idea that mPFC activity near the time of encoding potentiates subsequent consolidation processes, which together facilitate memory for negative information (Payne & Kensinger, 2010, 2011, 2018).

The results reported here also partially converge with recent work suggesting a causal role for the mPFC in gist-based false memories (Bovy et al., 2020; for related findings, see Berkers et al., 2017). Bovy and colleagues (2020) examined the relationship between gist-based emotional false memories and mPFC by stimulating the mPFC using an inhibitory TMS protocol, specifically continuous theta-burst stimulation (cTBS). Prior to encoding in an emotionally-valenced Deese-Roediger-McDermott paradigm, participants underwent a negative mood induction and received TMS to the mPFC with either a cTBS or control (i.e., 5Hz rTMS) protocol. They found that cTBS to mPFC during encoding reduced false recognition of negative critical lures on a test after a ~24-hour delay (including a night of sleep). The authors took these findings as evidence that the mPFC plays an important role in extracting schematic (or gist) information during encoding. Our findings extend the results described above. Here, we observed that sTBS – a TMS protocol thought to be associated with increased excitation (“long-term potentiation”, Wischnewski & Schutter, 2015) of the stimulated region – delivered to the mPFC during encoding facilitated memory for negative objects on a 24-hour delayed test. Moreover, we demonstrate that mPFC sTBS benefits emotional episodic memory by using a memory task consisting of complex emotional scenes as opposed to semantically related word lists. Most importantly, in addition to the 24-hour delayed test, we also included a short (30 minute) delayed memory test to better establish that mPFC activity during encoding interacted with subsequent consolidation processes. We observed that mPFC sTBS at encoding selectively enhanced negative memory only on the 24-hour delayed memory test. Our findings suggest that mPFC activity near the time of encoding interacts with downstream consolidation processes to support retention of emotional memories.

The present results also provide support for theories of the selective consolidation of emotional information (Kim & Payne, 2020; Payne & Kensinger, 2018; Richter-Levin & Akirav, 2003). Specifically, stimulation (i.e., neuromodulation) of the dorsal mPFC during encoding may have affected subsequent memory performance by upregulating regions important for emotional memory encoding and retrieval, including the ventral mPFC, amygdala and hippocampus (Bennion et al., 2015; Kensinger & Corkin, 2004; Murty et al., 2010; Payne & Kensinger, 2010). Future work could directly test this possibility by examining the effects of TMS in conjunction with fMRI measures of activity and connectivity.

There are some limitations of the present study that warrant mention. First, sTBS was interspersed throughout the encoding task to attempt to modulate brain regions, specifically the MTL, that are functionally connected with the mPFC and critically involved in the encoding of emotional information. Although we aimed to modulate mPFC-MTL activity during the encoding phase it is possible that our stimulation procedure may have also altered neural activity during early consolidation and/or retrieval processes (i.e., during the short delay recognition test) due to potentially long-lasting effects that extend 20-60 minutes following stimulation. Even though we are unaware of any evidence to suggest that the modified sTBS protocol implemented in the present study would have modulated neural activity beyond the encoding phase (e.g., modulating retrieval related processes), different stimulation protocols may provide further insights into the role of the mPFC activity near the time of encoding in selectively preserving negative information following a sleep filled delay. Furthermore, it is unclear if our results would differ had we used a different TMS protocol. For example, as described above, Bovy and colleagues found a causal role of the mPFC in gist-based false memories implementing cTBS while our sTBS protocol revealed a causal role of the mPFC in emotional memory (collapsed across specific and gist-based memories). Of relevance to this point, a recent meta-analysis of TMS effects on episodic memory found that offline 1 Hz rTMS led to larger enhancing effects on episodic memory compared to other stimulation protocols, including iTBS (Yeh & Rose, 2019). In addition, our excitatory assumption was based on the study by Demeters and colleagues (2016) that applied a similar sTBS protocol to the DLPFC and found enhanced word recognition. Coupled with our own enhanced memory for negative objects at a long delay results, we find it likely that sTBS had excitatory effects. Relatedly, if sTBS had inhibitory effects (like cTBS), then mPFC stimulation would have been expected to reduce memory performance (as observed by Bovy and colleagues, 2020), which we did not find. Our assumption that sTBS has similar effects to iTBS protocols despite the differences in the stimulation parameters (e.g., timing) is a limitation that is present across the literature. For example, the excitatory effects of iTBS are primarily based on stimulating the motor cortex but are often assumed to have similar effects when stimulating other cortical regions. Future work will be needed to probe if sTBS has similar excitatory effects as iTBS protocols.

Second, we only examined the effects of mPFC stimulation during memory encoding. It is possible that mPFC stimulation during a different stage of memory, such as during post-encoding periods, would lead to a different pattern of results. To draw stronger causal claims about the role of mPFC in emotional memory, we used an active control stimulation site (left MC) instead of another common control site (i.e., vertex) or sham stimulation. Prior findings have suggested vertex stimulation may modulate activity in the default mode network (Jung et al., 2016), which is involved in emotion (Sheline et al., 2009) and memory processes (Rugg & Vilberg, 2013). Although we cannot rule out the possibility that MC stimulation contributed to the effects on memory by impairing recognition of negative objects, we find this possibility unlikely for two reasons. First, memory performance in the MC stimulation group reported here was comparable to memory performance in a no-stimulation control group in a pilot study (not reported). Second, and most importantly, MC is not associated with emotional memory. Lastly, future work is needed to determine if different stimulation protocols (e.g., frequency, intensity, control stimulation site, stage of memory) will contribute to further insights about the role of encoding-consolidation interactions involving the mPFC and its associated network.

Future work should conduct a more direct investigation of the relationship between mPFC activity during encoding and sleep-based consolidation in emotional memory. Sleep affords an ideal environment for offline consolidation to transform, integrate, and preserve salient and future relevant information (Payne, 2011). Synchronous neural oscillations during sleep (e.g., theta oscillations, slow oscillations, spindles) have been observed in the amygdala, hippocampus, and prefrontal cortex, and may facilitate synaptic plasticity and selective memory consolidation during sleep (Kim & Payne, 2020; Rasch & Born, 2013). Examining the macro and microarchitecture of sleep in conjunction with causal methods like TMS will provide critical insight into the neural mechanisms of emotional memory encoding and consolidation.

In conclusion, the results of this study provide preliminary evidence for a causal role of the mPFC in selectively preserving negative memories. The current work moves beyond correlational findings and provides initial causal support for theories suggesting that activity and connectivity in an emotional memory network (e.g., mPFC-MTL) near the time of encoding interact with subsequent consolidation processes (perhaps during sleep) to form long-lasting emotional memories (Kim & Payne, 2020; Payne & Kensinger, 2018).

## Acknowledgments

Preparation of this manuscript was supported by the National Science Foundation (Grant BCS-1539361 awarded to J.D.P and E.A.K., and CAREER Grant 1848440 awarded to N.S.R) and from the University of Notre Dame William P. and Hazel B. Collegiate Chair funds (awarded to N.S.R). The authors would like to thank Katie McGuckin, Willian De Faria, Clare Hannon, Shannon O’Hara, and Frank Calabrese for contributing to data collection and analysis.

## References

Ballarini, F., Martínez, M. C., Díaz Perez, M., Moncada, D., & Viola, H. (2013). Memory in elementary school children is improved by an unrelated novel experience. PloS one, 8(6), e66875. https://doi.org/10.1371/journal.pone.0066875

Bennion, K. A., Mickley Steinmetz, K. R., Kensinger, E. A., & Payne, J. D. (2015). Sleep and cortisol interact to support memory consolidation. Cerebral Cortex, 25(3), 646–657. https://doi.org/10.1093/cercor/bht255

Berkers, R. M. W. J., Klumpers, F., & Fernández, G. (2016). Medial prefrontal–hippocampal connectivity during emotional memory encoding predicts individual differences in the loss of associative memory specificity. Neurobiology of Learning and Memory, 134, 44–54. https://doi.org/10/ggk792

Berkers, R. M. W. J., van der Linden, M., de Almeida, R. F., Müller, N. C. J., Bovy, L., Dresler, M., Morris, R. G. M., & Fernández, G. (2017). Transient medial prefrontal perturbation reduces false memory formation. Cortex, 88, 42–52. https://doi.org/10.1016/j.cortex.2016.12.015

Bovy, L., Berkers, R. M. W. J., Pottkämper, J. C. M., Varatheeswaran, R., Fernández, G., Tendolkar, I., & Dresler, M. (2020). Transcranial magnetic stimulation of the medial prefrontal cortex decreases emotional memory schemas. Cerebral Cortex, bhz329. https://doi.org/10.1093/cercor/bhz329

Daskalakis, Z. J., Farzan, F., Barr, M. S., Maller, J. J., Chen, R., & Fitzgerald, P. B. (2008). Long-interval cortical inhibition from the dorsolateral prefrontal cortex: A TMS-EEG study. Neuropsychopharmacology, 33(12), 2860–2869. https://doi.org/10.1038/npp.2008.22

Demeter, E., Mirdamadi, J. L., Meehan, S. K., & Taylor, S. F. (2016). Short theta burst stimulation to left frontal cortex prior to encoding enhances subsequent recognition memory. Cognitive, Affective, & Behavioral Neuroscience, 16(4), 724–735. https://doi.org/10.3758/s13415-016-0426-3

Dolcos, F., LaBar, K. S., & Cabeza, R. (2004). Dissociable effects of arousal and valence on prefrontal activity indexing emotional evaluation and subsequent memory: An event-related fMRI study. NeuroImage, 23(1), 64–74. https://doi.org/10/ckkrtb

Dunsmoor, J. E., Murty, V. P., Davachi, L., & Phelps, E. A. (2015). Emotional learning selectively and retroactively strengthens memories for related events. Nature, 520(7547), 345–348. https://doi.org/10.1038/nature14106

Fecchio, M., Pigorini, A., Comanducci, A., Sarasso, S., Casarotto, S., Premoli, I., Derchi, C.-C., Mazza, A., Russo, S., Resta, F., Ferrarelli, F., Mariotti, M., Ziemann, U., Massimini, M., & Rosanova, M. (2017). The spectral features of EEG responses to transcranial magnetic stimulation of the primary motor cortex depend on the amplitude of the motor evoked potentials. PloS One, 12(9), e0184910. https://doi.org/10.1371/journal.pone.0184910

Garoff, R. J., Slotnick, S. D., & Schacter, D. L. (2005). The neural origins of specific and general memory: The role of the fusiform cortex. Neuropsychologia, 43(6), 847–859. https://doi.org/10.1016/j.neuropsychologia.2004.09.014

Goto, Y., & Grace, A. A. (2008). Limbic and cortical information processing in the nucleus accumbens. Trends in Neurosciences, 31(11), 552–558. https://doi.org/10/b4tx84

Hermiller, M. S., Chen, Y. F., Parrish, T. B., & Voss, J. L. (2020). Evidence for immediate enhancement of hippocampal memory encoding by network-targeted theta-burst stimulation during concurrent fMRI. Journal of Neuroscience, 40(37), 7155–7168.

Hu, P., Stylos-Allan, M., & Walker, M. P. (2006). Sleep facilitates consolidation of emotional declarative memory. Psychological Science, 17(10), 891–898. https://doi.org/10/dkqxs9

Huang, Y.-Z., Edwards, M. J., Rounis, E., Bhatia, K. P., & Rothwell, J. C. (2005). Theta burst stimulation of the human motor cortex. Neuron, 45(2), 201–206. https://doi.org/10/d2gpww

Jung, J., Bungert, A., Bowtell, R., & Jackson, S. R. (2016). Vertex stimulation as a control site for transcranial magnetic stimulation: A Concurrent TMS/fMRI Study. Brain Stimulation, 9(1), 58–64. https://doi.org/10.1016/j.brs.2015.09.008

Kensinger, E. A., & Corkin, S. (2004). Two routes to emotional memory: Distinct neural processes for valence and arousal. Proceedings of the National Academy of Sciences of the United States of America, 101(9), 3310–3315. https://doi.org/10/ftzx6z

Kensinger, E. A., Garoff-Eaton, R. J., & Schacter, D. L. (2007a). Effects of emotion on memory specificity: Memory trade-offs elicited by negative visually arousing stimuli. Journal of Memory and Language, 56(4), 575–591. https://doi.org/10.1016/j.jml.2006.05.004

Kensinger, E. A., Garoff-Eaton, R. J., & Schacter, D. L. (2007b). How negative emotion enhances the visual specificity of a memory. Journal of Cognitive Neuroscience, 19(11), 1872–1887. https://doi.org/10.1162/jocn.2007.19.11.1872

Kim, S. Y., & Payne, J. D. (2020). Neural correlates of sleep, stress, and selective memory consolidation. Current Opinion in Behavioral Sciences, 33, 57–64. https://doi.org/10/ggmtgv

Lenth, R. (2020). Emmeans: Estimated marginal means, aka least-squares means. Retrieved from https://CRAN.R-project.org/package=emmeans

Murty, V. P., Ritchey, M., Adcock, R. A., & LaBar, K. S. (2010). fMRI studies of successful emotional memory encoding: A quantitative meta-analysis. Neuropsychologia, 48(12), 3459–3469. https://doi.org/10/dwkvv8

Okamoto, M., Dan, H., Sakamoto, K., Takeo, K., Shimizu, K., Kohno, S., Oda, I., Isobe, S., Suzuki, T., Kohyama, K., & Dan, I. (2004). Three-dimensional probabilistic anatomical cranio-cerebral correlation via the international 10–20 system oriented for transcranial functional brain mapping. NeuroImage, 21(1), 99–111. https://doi.org/10.1016/j.neuroimage.2003.08.026

Okamoto, M., & Dan, I. (2005). Automated cortical projection of head-surface locations for transcranial functional brain mapping. NeuroImage, 26(1), 18–28. https://doi.org/10.1016/j.neuroimage.2005.01.018

Patil, A., Murty, V. P., Dunsmoor, J. E., Phelps, E. A., & Davachi, L. (2016). Reward retroactively enhances memory consolidation for related items. Learning & memory, 24(1), 65–69. https://doi.org/10.1101/lm.042978.116

Payne, J. D., Chambers, A. M., & Kensinger, E. A. (2012). Sleep promotes lasting changes in selective memory for emotional scenes. Frontiers in Integrative Neuroscience, 6. https://doi.org/10/ggk9f7

Payne, J. D., & Kensinger, E. A. (2010). Sleep’s role in the consolidation of emotional episodic memories. Current Directions in Psychological Science, 19(5), 290–295. https://doi.org/10/b6xkvt

Payne J. D. (2011). Sleep on it!: Stabilizing and transforming memories during sleep. Nature Neuroscience, 14(3), 272–274. https://doi.org/10.1038/nn0311-272

Payne, J. D., & Kensinger, E. A. (2011). Sleep leads to changes in the emotional memory trace: Evidence from fMRI. Journal of Cognitive Neuroscience, 23(6), 1285–1297. https://doi.org/10/btvfds

Payne, J. D., & Kensinger, E. A. (2018). Stress, sleep, and the selective consolidation of emotional memories. Current Opinion in Behavioral Sciences, 19, 36–43. https://doi.org/10/ggk8cj

Payne, J. D., Stickgold, R., Swanberg, K., & Kensinger, E. A. (2008). Sleep preferentially enhances memory for emotional components of scenes. Psychological Science, 19(8), 781–788. https://doi.org/10.1111/j.1467-9280.2008.02157.x

Rasch, B., & Born, J. (2013). About sleep’s role in memory. Physiological Reviews, 93(2), 681–766. https://doi.org/10.1152/physrev.00032.2012

Richter-Levin, G., & Akirav, I. (2003). Emotional tagging of memory formation—In the search for neural mechanisms. Brain Research Reviews, 43(3), 247–256. https://doi.org/10/c7v5xn

Rossi, S., Hallett, M., Rossini, P. M., & Pascual-Leone, A. (2009). Safety, ethical considerations, and application guidelines for the use of transcranial magnetic stimulation in clinical practice and research. Clinical Neurophysiology, 120(12), 2008–2039. https://doi.org/10/fwp29f

Rugg, M. D., & Vilberg, K. L. (2013). Brain networks underlying episodic memory retrieval. Current Opinion in Neurobiology, 23(2), 255–260. https://doi.org/10.1016/j.conb.2012.11.005

Sack, A. T., Cohen Kadosh, R., Schuhmann, T., Moerel, M., Walsh, V., & Goebel, R. (2009). Optimizing functional accuracy of TMS in cognitive studies: A comparison of methods. Journal of Cognitive Neuroscience, 21(2), 207–221. https://doi.org/10.1162/jocn.2009.21126

Sheline, Y. I., Barch, D. M., Price, J. L., Rundle, M. M., Vaishnavi, S. N., Snyder, A. Z., Mintun, M. A., Wang, S., Coalson, R. S., & Raichle, M. E. (2009). The default mode network and self-referential processes in depression. Proceedings of the National Academy of Sciences, 106(6), 1942–1947. https://doi.org/10.1073/pnas.0812686106

Singmann, H., Bolker, B., Westfall, J., Aust, F., & Ben-Shachar, M. S. (2020). Afex: Analysis of factorial experiments. Retrieved from https://CRAN.R-project.org/package=afex

Stanny, C. J., & Johnson, T. C. (2000). Effects of stress induced by a simulated shooting on recall by police and citizen witnesses. The American Journal of Psychology, 113(3), 359–386. https://doi.org/10/cmbhhf

Sterpenich, V., Albouy, G., Darsaud, A., Schmidt, C., Vandewalle, G., Dang Vu, T. T., Desseilles, M., Phillips, C., Degueldre, C., Balteau, E., Collette, F., Luxen, A., & Maquet, P. (2009). Sleep promotes the neural reorganization of remote emotional memory. The Journal of Neuroscience, 29(16), 5143–5152. https://doi.org/10/c3s6w3

Tang, Y., Jiao, X., Wang, J., Zhu, T., Zhou, J., Qian, Z., Zhang, T., Cui, H., Li, H., Tang, X., Xu, L., Zhang, L., Wei, Y., Sheng, J., Liu, L., & Wang, J. (2019). Dynamic functional connectivity within the fronto-limbic network induced by intermittent theta-burst stimulation: A pilot study. Frontiers in neuroscience, 13, 944. https://doi.org/10.3389/fnins.2019.00944

Tambini, A., Rimmele, U., Phelps, E. A., & Davachi, L. (2017). Emotional brain states carry over and enhance future memory formation. Nature neuroscience, 20(2), 271–278. https://doi.org/10.1038/nn.4468

Wang, J. X., Rogers, L. M., Gross, E. Z., Ryals, A. J., Dokucu, M. E., Brandstatt, K. L., Hermiller, M. S., & Voss, J. L. (2014). Targeted enhancement of cortical-hippocampal brain networks and associative memory. Science (New York, N.Y.), 345(6200), 1054–1057. https://doi.org/10.1126/science.1252900

Waring, J. D., Payne, J. D., Schacter, D. L., & Kensinger, E. A. (2010). Impact of individual differences upon emotion-induced memory trade-offs. Cognition & Emotion, 24(1), 150–167. https://doi.org/10.1080/02699930802618918

Warren, K. N., Hermiller, M. S., Nilakantan, A. S., & Voss, J. L. (2019). Stimulating the hippocampal posterior-medial network enhances task-dependent connectivity and memory. eLife, 8, e49458. https://doi.org/10.7554/eLife.49458

Wischnewski, M., & Schutter, D. J. (2015). Efficacy and time course of theta burst stimulation in healthy humans. Brain stimulation, 8(4), 685–692.

Yarkoni, T., Poldrack, R. A., Nichols, T. E., Van Essen, D. C., & Wager, T. D. (2011). Large-scale automated synthesis of human functional neuroimaging data. Nature methods, 8(8), 665–670.

Yeh, N., & Rose, N. S. (2019). How can transcranial magnetic stimulation be used to modulate episodic memory?: A systematic review and meta-analysis. Frontiers in Psychology, 10. https://doi.org/10.3389/fpsyg.2019.00993

Yonelinas, A. P., & Jacoby, L. L. (1995). The relation between remembering and knowing as bases for recognition: Effects of size congruency. Journal of Memory and Language, 34(5), 622–643. https://doi.org/10/b3gq9t

